# Kidney osteoclast factors and matrix metalloproteinase expression in a mice model of diet-induced obesity and diabetes

**DOI:** 10.1101/615716

**Authors:** Francine dos Santos-Macedo, Bianca Martins-Gregorio, Elan Cardozo Paes-de-Almeida, Leonardo de Souza Mendonça, Rebeca de Souza Azevedo, Caroline Fernandes-Santos

**Author notes:** Corresponding Author: Rua Dr. Silvio Henrique Braune, 22, Centro, Nova Friburgo, RJ, Brazil, 28.625-650.

## Abstract

The role of RANKL/RANK/OPG system on bone remodeling is well known, and there is evidence that it is also important to cardiovascular and kidney pathology, although the underlying mechanisms are not elucidated so far. Thus, we investigated in a mice model of diet-induced obesity and diabetes if renal histopathological changes are associated with the expression of RANKL/RANK/OPG system and matrix metalloproteinases (MMPs). Three months old C57BL/6 mice were fed with control (AIN93M) or high-fat high sucrose (HFHS) diets for 21 weeks (CEUA/UFF #647/15). The HFHS group showed weight gain (+35%, P=0.0001), increased epididymal, inguinal and retroperitoneal fat pad weight (+121 %, P = 0.0006; +287 %, P = 0.0007 and; +286 %, P < 0.0001, respectively), and hyperglycemia (+43%, P=0.02). The kidney of some HFHS fed mice displayed mononuclear inflammatory cell infiltrate (40%), perivascular fibrosis (20%), and focal tubule mineralization (20%). Glomeruli hypertrophy was not detected. Unexpectedly, OPG, RANK, MMP-2 and MMP-9 expression was not altered in HFHS groups (Western blot analysis). In conclusion, the expression of RANKL/RANK/OPG system proteins and MMPs was not influenced by diet-induced obesity and diabetes in the kidney of male C57BL/6 mice, although some adverse histopathological remodeling is noticed in the renal tissue.

## INTRODUCTION

Osteoprotegerin (OPG) is a soluble receptor that acts as a site competitor of the receptor activator of NF-κB ligand (RANKL) [1]. In turn, RANKL is the ligand of the **receptor activator of** NF-κB (RANK**) [2].** The biological effect of RANKL on bone reabsorption is produced when it binds to RANK on the surface of pre-osteoclasts, which results in osteoclast differentiation, activation, and survival. OPG competes with RANKL binding sites, neutralizing its effect and thus OPG was initially identified as a critical factor on bone remodeling [3, 4]. Although the role of RANKL/RANK/OPG system on bone remodeling is widely well-known, its function in cardiovascular and renal systems is yet under investigation.

The RANKL/RANK/OPG system has been associated with the development and progression of cardiovascular and kidney diseases since these proteins are also involved in the immune process [5, 6]. However, the evidence is limited to blood analysis and studies with tissue samples are scarce. Plasma OPG is increased in diabetic patients with nephropathy [7, 8], but the mechanism is not well understood. Nascimento et al. [9] hypothesize that the concentration of circulating OPG would be associated with the presence of atherosclerosis and that OPG would predict mortality in patients with advanced kidney disease. Also, the author reported that circulating OPG could be a useful biomarker for the clinical evaluation of patients suffering from nephropathies, and thus a possible target for therapeutic intervention.

Extracellular matrix (ECM) remodeling is a common adverse finding in several kidney diseases, and matrix metalloproteinases (MMPs) are a large family of proteases that play an essential role in this process since they degrade all types of ECM proteins [10, 11]. MMPs are important in the development and progression of chronic kidney disease, and MMP-2 and MMP-9 are prominent in this process since their expression are upregulated during renal fibrosis [12]. Immune system cells, such as Th17, seem to induce the expression of MMPs through activation by signaling proteins from the RANKL/RANK/OPG system, and this free activation is of great importance to the remodeling process [13].

Overall, obesity and diabetes are conditions that chronically leads to renal complication, such as glomerulonephritis, nephropathies, and nephrosclerosis [14]. Thus, it is of interest to investigate if kidney histopathological changes found in these conditions are associated with altered expression of the RANKL/RANK/OPG system and MMPs. To accomplish that, we used a mice model of diet-induced obesity and diabetes to investigate if renal histopathological finding correlates with the expression of RANKL/RANK/OPG system and MMPs.

## MATERIAL AND METHODS

### Ethical aspects

The local Ethics Committee approved the handling and experimental protocols to Care and Use of Laboratory Animals (CEUA#647/15). The study was performed following the Animal Research Reporting In vivo Experiments ARRIVE guidelines and the Guideline for the Care and Use of Laboratory Animals (US NIH Publication N° 85-23. Revised 1996) [15].

### Animal husbandry and diet

Mice were kept in the animal care facility of the Laboratorio Multiusuario de Pesquisa Biomedica, Universidade Federal Fluminense, Nova Friburgo, Rio de Janeiro, Brazil. Twelve male C57BL/6 mice at three months old were maintained in collective polycarbonate microisolators (six mice per cage) of 30×20×23/28×18×11 cm external/internal dimensions, with a wire bar lid that serves as food hopper and water bottle holder on ventilated racks (Scienlabor Industrial Equipment, SP, Brazil). The housing conditions were 12 h light/dark cycles, 21±2°C, 60±10% humidity and air exhaustion cycle of 15 min/h. Mice were feed for twenty-one weeks with a purified AIN93M diet [16] (control group, C), or a high-fat high-sucrose diet (HFHS group) modified from the AIN93M diet (Pragsolucoes, Jau, Sao Paulo, Brazil) to induce obesity and diabetes [17-19]. Briefly, the AIN93M diet had 3.81 kcal/g and consisted of 15% protein (casein), 9% fat (soybean oil), and 76% carbohydrate (65% as corn starch and 11% as sucrose). The HFHS diet had 4.71 kcal/g and consisted of 14% protein (casein), 42% fat (9% as soybean oil and 33% as lard), and 44% carbohydrate (19% as corn starch and 25% as sucrose). Mice had ad libitum access to water and food. Daily food intake and energy, as well as weekly body mass (BM), were recorded.

### Blood glucose and triacylglycerol

Blood glucose before and after 21 weeks of diet intake was assessed by performing a small incision at the tip of the animal tail for blood collection after six hours of fasting (Accu-Check Performa glucose meter, Roche). Glucose intolerance was assessed after 21 weeks on diet by the oral glucose tolerance test (OGTT). Animals received a 50 % glucose solution by orogastric gavage after six hours of fasting (2 g/kg). Blood was also collected through a small incision at the tail tip before glucose administration and 15, 30, 60, 90 and 120 minutes after glucose gavage. The area under the curve (AUC) was calculated to assess glucose intolerance [20]. At the end of the experiment, mice were anesthetized with ketamine (100 mg/kg) and xylazine (10 mg/kg) ip. Blood was withdrawn from the heart (right atrium) and allowed to retract at room temperature for half an hour. After centrifugation, serum was isolated and used for triacylglycerol assay (colorimetric assay, TGML-0517, ELITech Clinical Systems).

### Kidney histopathology

At euthanasia, both kidneys were removed and weighed, as well as visceral (epididymal and retroperitoneal) and subcutaneous (inguinal) white fat pads. The left kidney of six mice per group was used for histopathology analysis. They were cut on their longitudinal axis, fixed for 48 hours in 4% buffered formalin pH 7.2, underwent routine histological processing, and were embedded in Paraplast plus (P3683, Sigma). Then, non-consecutive 3 μm thick slices were stained with hematoxylin and eosin or Masson’s trichrome.

For quantitative analysis, six digital images per mice stained with hematoxylin and eosin were acquired at 40x magnification (Evos XL, ThermoScientific). To assess glomeruli diameter, at least thirty glomeruli had their diameter measured at two perpendicular axes (ImageJ, NIH, https://imagej.nih.gov/ij/). Kidney sections stained with Masson’s trichrome were used to qualitatively evaluate the presence of fibrosis, inflammation, hyalinosis, and mineralization at 20x magnification (Laser microdissection with PALMBeam microscope, Zeiss). These findings were classified as focal (isolated in a single spot), multifocal (located at several regions), interstitial (at the extracellular matrix around tubules) or perivascular (around blood vessels).

### Western blot

Samples of the right kidney were homogenized in RIPA buffer (Ika T10 Basic Ultra-Turrax®) and centrifuged. The supernatant was collected for protein quantification (Pierce BCA protein assay kit, code 23225, Thermo Fisher Scientific), 10 µg of proteins were separated by polyacrylamide gel electrophoresis (SDS-PAGE), and then wet transferred (Mini VE Complete, Pharmacia Biotech) from the gel to a polyvinylidene fluoride (PVDF) membrane (Amersham™ Hybond® P). For protein detection, PVDF membranes were blocked with 5% non-fat dry milk in TTBS for 1 h, then incubated 2h at room temperature and then overnight 4°C with primary antibodies (1:500, anti-OPG sc-8468, Lot#L1715; 1:500, anti-RANK, sc-9072, Lot # G0715; 1:1,000, anti-MMP-2, sc-53630; 1:1,000, anti-MMP-9, sc-21733 – Santa Cruz Biotechnology) or beta-actin as endogenous control (1:500, B1305554-40TST, SG100806AD, Sigma Aldrich). On the next day, after 1 h at room temperature, PVDF membranes were incubated with secondary antibodies for 1 h (1:1,000, anti-rabbit, sc-2004, Lot#I0513; 1:5,000, anti-goat, sc-2020, Lot#H2113; 1:5,000, anti-mouse, sc-2005 Lot#B2014 – Santa Cruz Biotechnology). The antigen-antibody reaction was detected by ECL incubation (GE Healthcare Life Sciences) for 5 minutes on Licor C-DiGit Blot Scanner. Membranes were stripped with 2% glycine pH 2.0 for reuse. Protein band density was analyzed using the Gel Analyzer feature of ImageJ.

### Statistical analysis

Data are expressed as mean ± SD, and they were tested for normality and homoscedasticity of variances. Comparison among groups was made by Student’s t-test, except for food and energy intake (Mann Whitney’s test). All analysis considered a P-value of 0.05 (Graph Pad® Prism v.6.0, La Jolla, CA, USA).

## RESULTS

### Obesity and diabetes in C57BL/6 mice fed an HFHS diet

Mice fed with the HFHS diet showed a significant increase of BM after ten weeks of HFHS feeding (Figure 1a). After 21 weeks, BM increased by 118% (P=0.0007, Figure 1b). Compared with C group, HFHS fed mice ingested less food (−20%, P<0.0001) but energy intake remained similar between groups (Figure 1c-d). BM gain was followed by increased body adiposity since epididymal, inguinal and retroperitoneal WAT pads were consistently higher in HFHS group mice than C group (+121%, P=0.0006; +287%, P=0.0007 and; +286%, P<0.0001, respectively, Figure 1e-g). Obesity was followed by hyperglycemia and glucose intolerance in the HFHS group (Figure 2). After 21 weeks, fasting glucose increased by 51 % (P=0.0022, Figure 2c), and the area under the curve of OGTT increased by 31% (P=0.02, Figure 2d). Serum triacylglycerol was not affected by the HFHS diet (Figure 2e).

**Figure 1.**
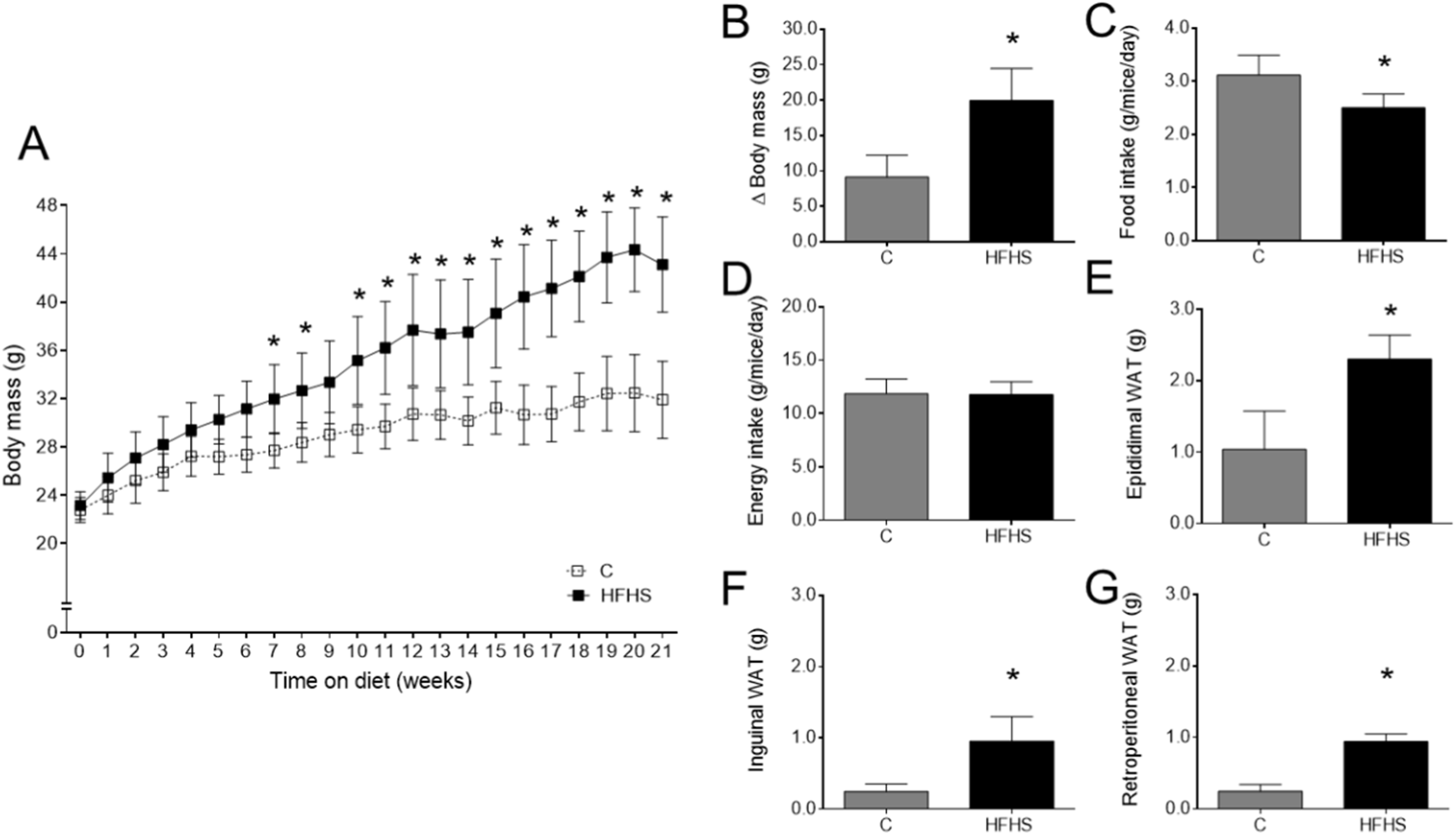
Mice fed with high-fat high sucrose (HFHS) diet for 21 weeks have increased body mass (a-b). Food intake was smaller in HFHS compared to control (C) group, but energy intake remained similar (c-d). All white fat depots analyzed were heavier in HFHS mice compared with C group after 21 weeks of diet feeding (e-g). N = 6/ group, mean ± SD, Student’s t-test, P < 0.05 vs. C group.

**Figure 2.**
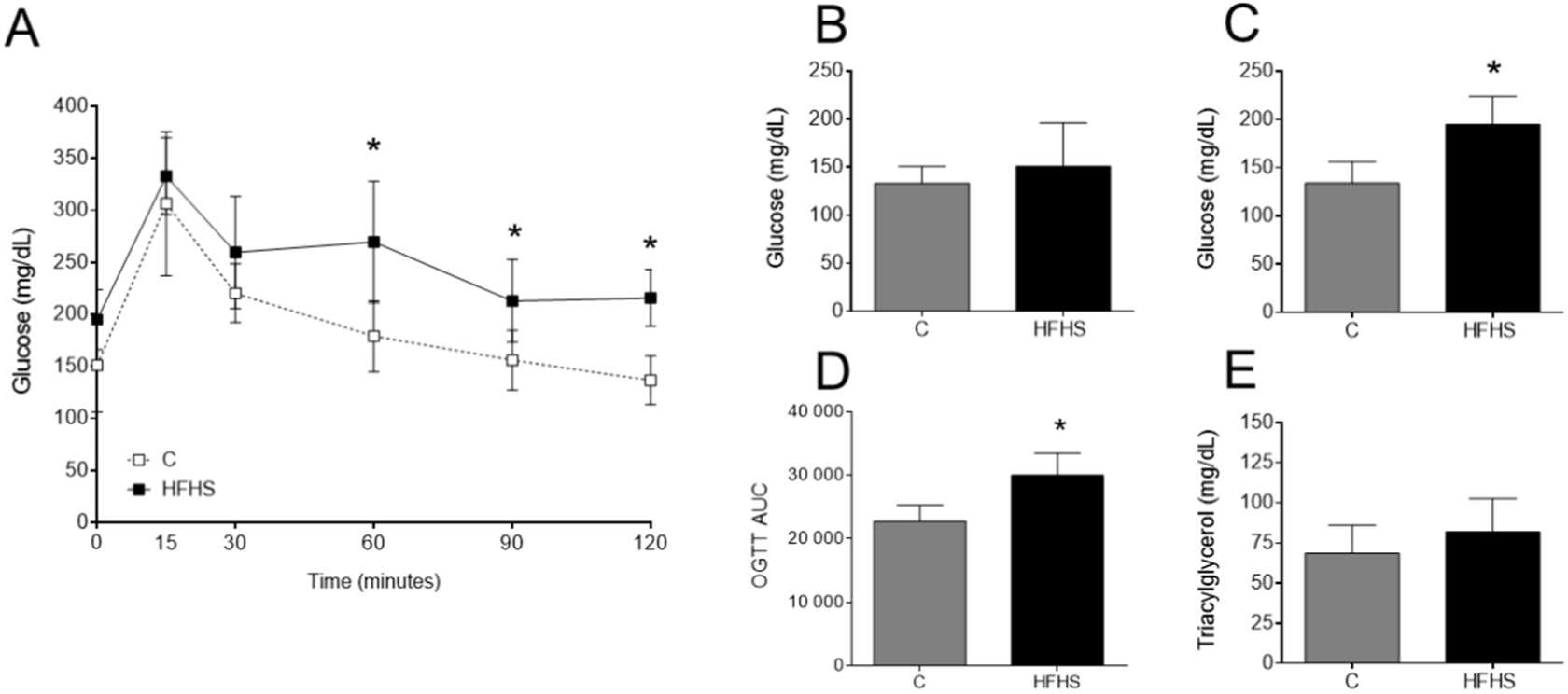
Glucose metabolism. Fasting glucose before diet administration is shown in (b) and 21 weeks after diet feeding is shown in (c). Glucose intolerance was evaluated by the oral glucose tolerance test (OGTT) curve (a) and its area under the curve (d). Serum triacylglycerol is shown in (e). N = 6/ group, mean ± SD, Student’s t-test, P < 0.05 vs. C group.

### Kidney histopathology and protein expression

Kidney weight and average glomeruli diameter did now change by chronic HFHS diet intake (Figure 3a-b). However, qualitative histopathology showed many infiltrates of mononuclear inflammatory cells in the HFHS group (50% of mice), and it was often seen associated with interstitial fibrosis (Figure 3c). Perivascular fibrosis and focal tubule mineralization were seen only in the HFHS group. Hyalinosis was found in both C and HFHS groups (Figure 3c). Red arrows in Figure 3d-h exemplifies the main histopathological findings displayed in Figure 3c. Unexpectedly, no change was detected in OPG, RANK, MMP-2 and MMP-9 protein expression in the kidney of mice fed during 21 weeks with the HFHS diet (Figure 4).

**Figure 3.**
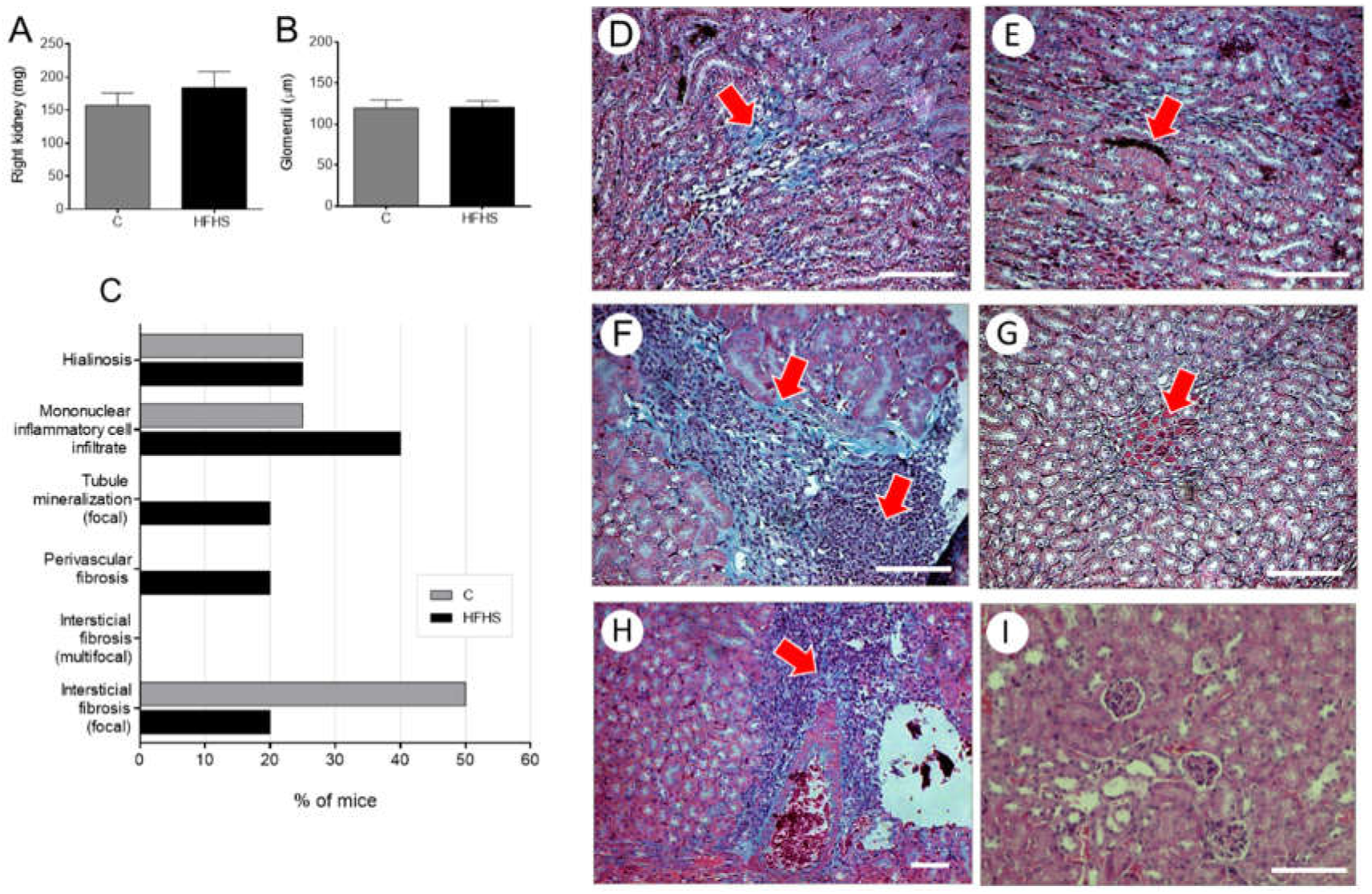
Kidney weight (a) and glomeruli size (b) were not changed after 21 weeks on a diet rich in fat and sucrose. Histopathologic features evaluated are shown in (c), as a percentage of mice per group that displayed each feature. The photomicrographs (d-i) illustrate the histopathological findings in HFHS group (red arrows): mononuclear inflammatory infiltrate associated with discrete fibrosis (d), mineralization (e), mononuclear inflammatory infiltrate associated with focal fibrosis (f), hyalinosis (g), perivascular fibrosis (adventitia) associated with mononuclear inflammatory infiltrate (h) and normal aspect of renal cortex with some glomeruli (i). N = 6/ group, mean ± SD, Student’s t-test, P < 0.05 vs. C group.

**Figure 4.**
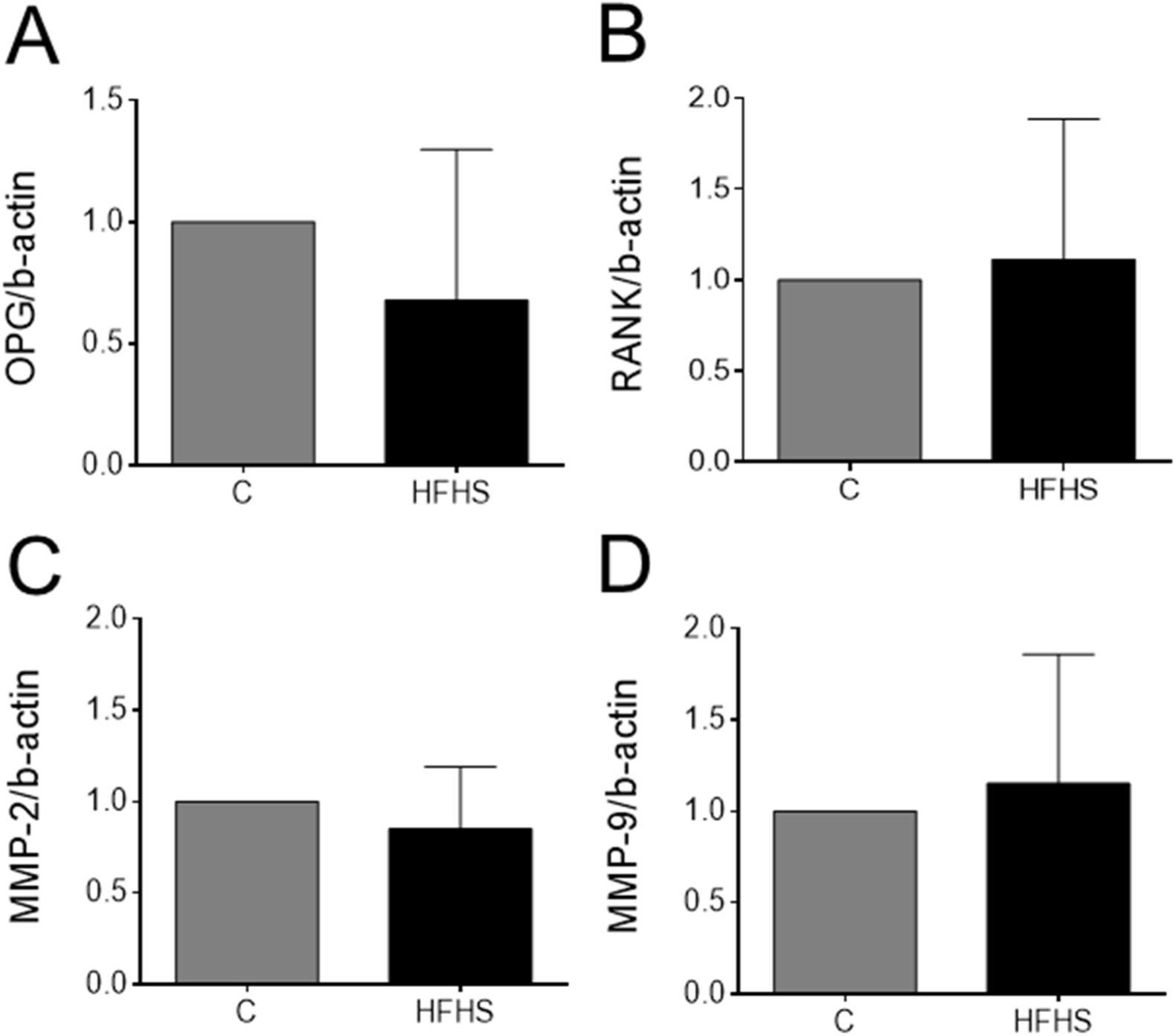
– Osteoclast factors and matrix metalloproteinase expression by Western blot. Osteoprotegerin (OPG) (a), receptor activator of NF-κB (RANK) (b), matrix metalloproteinase (MMP)-2 (c) and MMP-9 (d). N = 6/ group, mean ± SD, Student’s t-test, P < 0.05 vs. C group.

## DISCUSSION

In the present study, C57BL/6 mice were fed with a HFHS diet to induce overweight/ obesity, diabetes, and histopathological kidney injuries. The HFHS diet was offered for 21 weeks because other studies have already shown that this time is required to promote kidney injury [21, 22]. In our study, the HFHS group had kidney fibrosis, mineralization, and hyalinosis, but glomeruli hypertrophy was not seen. Although some histopathological changes were also found in the C group, they were already expected in some healthy mice from the colony, as reported by Scudamore et al. (e.g., inflammatory cell infiltrate and foci in male mice) [23]. Unexpectedly, modulation of RANKL/RANK/OPG system proteins, as well as MMP-2 and MMP-9 enzymes, were not detected by the current protocol of chronic diet-induced obesity and diabetes. Also, we could not find other reports in the literature that investigate these same proteins in similar conditions that could be confronted with our data.

A study with diabetic patients with nephropathy observed that plasma OPG concentration was elevated, suggesting that it would reflect renal injury because OPG elevation is associated with microvascular injury and endothelial dysfunction [7]. Since nephropathy is a microvascular complication, plasma OPG concentration would reflect microvascular dysfunction. They also investigated renal biopsy, and found no glomeruli OPG stain, unlike tubular cells [6]. Authors say that OPG would be more expressed in tubular cells compared with glomeruli and that Western blot would be a more suitable technique to investigate OPG expression in the renal tissue. In the present study in a mice model of diet-induced obesity and diabetes, OPG expression was investigated by Western blot, and we could not find changes in its expression compared with control mice.

There is a need for further researches to fully define the therapeutic potential of the RANKL/RANK/OPG system in patients with renal injury, because it is thought that OPG can be a biomarker in patients with renal disease, predicting vascular calcification and survival. For instance, a study with mice lacking OPG observed that it prevents smooth muscle cells calcification and protects renal cells from TRAIL-induced death. OPG can then be a biomarker in patients with renal disease, predicting vascular calcification and survival [24].

Kidney fibrosis is almost always preceded by and closely associated with inflammation [25, 26], since the aim of the inflammatory process is to eliminate the original insult, by removing cells and ECM debris. Inflammatory cells infiltrate in the tissue, fibroblasts are activated, and there is excessive deposition of ECM components leading to fibrosis [27]. Under normal conditions, the human kidney produces MMP-2 and MMP-9, always at a low level [28]. During the fibrosis process, those MMPs are rapidly upregulated, and their abnormal expression can be critical to the development of glomerulosclerosis [28, 29]. In the current study, we observed the inflammatory infiltrate often associated with fibrosis as expected, but when the whole kidney homogenate is analyzed, we could not detect changes in MMP-2 and MMP-9 expression. One hypothesis is that histopathological changes were mainly focal and not all mice were affected for the time length of HFHS diet administration, thus average MMPs expression is undetectable (if present).

Studies in mice knockout to MMP-2 and MMP-9 have helped to elucidate their role on kidney fibrosis. MMP-2^-/-^ mice with streptozotocin-induced diabetic nephropathy have increased renal fibrosis due to impaired collagen degradation since this MMP is not a collagenase and would not directly degrade this component [30]. Another study investigated the progression of renal interstitial fibrosis by a unilateral ureteral obstruction in MMP-9^−/−^ mice, and they reported reduced kidney fibrosis, suggesting that MMP-9 is a pro-fibrotic mediator [6, 31].

Although it is not yet clear whether the increase of RANKL/RANK/OPG system proteins is beneficial to the body, some authors believe that the system, in particular, OPG, exerts a protective role by both preventing cells from calcification and death inducers [4]. To date, our study initiated an investigation about the role of the RANKL/RANK/OPG system in diet-induced obesity and diabetes in mice, a field not exploited so far. Even so, further studies are still necessary to elucidate the role of the system in kidney histopathological changes associated with obesity and diabetes.

## CONCLUSION

In conclusion, the expression of RANKL/RANK/OPG system proteins and MMPs is not influenced by diet-induced obesity and diabetes in the kidney of male C57BL/6 mice, although some adverse histopathological remodeling is noticed in the renal tissue.

## Acknowledgments

Authors are thankful to Ana Luiza Martins Gallo e Joao Augusto Mulin Montechiari Machado for their help in animal care, and to Dilliane da Paixao Rodrigues Almeida for her technical assistance in sample preparation for histopathology analysis.

## Conflict of interest

The authors declare that they have no conflict of interests.

## Funding

This study was supported by a grant from PROPPI (Fluminense Federal University). Student’s scholarship was provided by CAPES (Coordination for the Improvement of Higher Education Personnel – Coordenação de Aperfeiçoamento de Pessoal de Nível Superior – Brasil/ Finance Code 001).

## Authorship

F.S-M performed the experiment, collected, analyzed, and interpreted data, and wrote the manuscript draft. B.M.G. and E.C.P-A. assisted the histopathology analysis. C.F-S designed the experiment, interpreted data, and wrote the manuscript draft. L.S.M. and R.S.A designed the experiment and gathered a grant for the research. All authors read and approved the submitted and final manuscript versions.

## Availability of data and material

The datasets used and analyzed during the current study are available from the corresponding author on reasonable request.

## REFERENCES

[1] Khosla, S., et al., Correlates of osteoprotegerin levels in women and men. Osteoporos Int, 2002. 13(5): p. 394–9.

[2] yrovola, J.B., et al., Root resorption and the OPG/RANKL/RANK system: a mini review. J Oral Sci, 2008. 50(4): p. 367–76.

[3] Hofbauer, L.C., Pathophysiology of RANK ligand (RANKL) and osteoprotegerin (OPG). Ann Endocrinol (Paris), 2006. 67(2): p. 139–41.

[4] Malliga, D.E., D. Wagner, and A. Fahrleitner-Pammer The role of osteoprotegerin (OPG) receptor activator for nuclear factor kappaB ligand (RANKL) in cardiovascular pathology – a review. 2011.

[5] Bernardi, S., et al., Osteoprotegerin increases in metabolic syndrome and promotes adipose tissue proinflammatory changes. Mol Cell Endocrinol, 2014. 394(1-2): p. 13–20.

[6] Wang, S.T., et al., The plasma osteoprotegerin level and osteoprotegerin expression in renal biopsy tissue are increased in type 2 diabetes with nephropathy. Exp Clin Endocrinol Diabetes, 2015. 123(2): p. 106–11.

[7] Wang, S.T., et al., Increased plasma osteoprotegerin concentrations in Type 1 diabetes with albuminuria. Clin Nephrol, 2013. 79(3): p. 192–8.

[8] Xiang, G.D., et al., Association between plasma osteoprotegerin concentrations and urinary albumin excretion in Type 2 diabetes. Diabet Med, 2009. 26(4): p. 397–403.

[9] Nascimento, M.M., et al., Elevated levels of plasma osteoprotegerin are associated with allcause mortality risk and atherosclerosis in patients with stages 3 to 5 chronic kidney disease. Braz J Med Biol Res, 2014. 47(11): p. 995–1002.

[10] Gargiulo, S., et al., Metalloproteinases and metalloproteinase inhibitors in age-related diseases. Curr Pharm Des, 2014. 20(18): p. 2993–3018.

[11] Amin, M., et al., Regulation and involvement of matrix metalloproteinases in vascular diseases. Front Biosci (Landmark Ed), 2016. 21: p. 89–118.

[12] Cheng, Z., et al., MMP-2 and 9 in Chronic Kidney Disease. Int J Mol Sci, 2017. 18(4).

[13] Liu, W., et al., Osteoprotegerin/RANK/RANKL axis in cardiac remodeling due to immunoinflammatory myocardial disease. Exp Mol Pathol, 2008. 84(3): p. 213–7.

[14] Kopple, J.D., Obesity and chronic kidney disease. J Ren Nutr, 2010. 20(5 Suppl): p. S29–30.

[15] Kilkenny, C., et al., Improving bioscience research reporting: The ARRIVE guidelines for reporting animal research. J Pharmacol Pharmacother, 2010. 1(2): p. 94–9.

[16] Reeves, P.G., F.H. Nielsen, and G.C. Fahey, Jr., AIN-93 purified diets for laboratory rodents: final report of the American Institute of Nutrition ad hoc writing committee on the reformulation of the AIN-76A rodent diet. J Nutr, 1993. 123(11): p. 1939–51.

[17] Surwit, R.S., et al., Differential effects of fat and sucrose on the development of obesity and diabetes in C57BL/6J and A/J mice. Metabolism, 1995. 44(5): p. 645–51.

[18] Fernandes-Santos, C., et al., Pan-PPAR agonist beneficial effects in overweight mice fed a high-fat high-sucrose diet. Nutrition, 2009. 25(7-8): p. 818–27.

[19] Fernandes-Santos, C., et al., Rosiglitazone aggravates nonalcoholic Fatty pancreatic disease in C57BL/6 mice fed high-fat and high-sucrose diet. Pancreas, 2009. 38(3): p. e80–6.

[20] Andrikopoulos, S., et al., Evaluating the glucose tolerance test in mice. Am J Physiol Endocrinol Metab, 2008. 295(6): p. E1323–32.

[21] Wei, P., et al., Glomerular structural and functional changes in a high-fat diet mouse model of early-stage Type 2 diabetes. Diabetologia, 2004. 47(9): p. 1541–9.

[22] Tain, Y.L., et al., High Fat Diets Sex-Specifically Affect the Renal Transcriptome and Program Obesity, Kidney Injury, and Hypertension in the Offspring. Nutrients, 2017. 9(4).

[23] Scudamore, C., 31/05/2018 A Practical Guide to the Histology of the Mouse Wiley Online Books https://onlinelibrary.wiley.com/doi/book/10.1002/97811187895681/4 Wiley Online Books A practical guide to the histology of the mouse. 2014.

[24] Montañez-Barragán, A., et al., Osteoprotegerin and kidney disease. J Nephrol, 2014. 27(6): p. 607–17.

[25] Krensky, A.M. and Y.T. Ahn, Mechanisms of disease: regulation of RANTES (CCL5) in renal disease. Nat Clin Pract Nephrol, 2007. 3(3): p. 164–70.

[26] Lange-Sperandio, B., et al., Leukocytes induce epithelial to mesenchymal transition after unilateral ureteral obstruction in neonatal mice. Am J Pathol, 2007. 171(3): p. 861–71.

[27] Lee, S.B. and R. Kalluri, Mechanistic connection between inflammation and fibrosis. Kidney Int Suppl, 2010(119): p. S22–6.

[28] Wiercinska, E., et al., The TGF-β/Smad pathway induces breast cancer cell invasion through the up-regulation of matrix metalloproteinase 2 and 9 in a spheroid invasion model system. Breast Cancer Res Treat, 2011. 128(3): p. 657–66.

[29] Meran, S. and R. Steadman, Fibroblasts and myofibroblasts in renal fibrosis. Int J Exp Pathol, 2011. 92(3): p. 158–67.

[30] Takamiya, Y., et al., Experimental diabetic nephropathy is accelerated in matrix metalloproteinase-2 knockout mice. Nephrol Dial Transplant, 2013. 28(1): p. 55–62.

[31] Wang, X., et al., Mice lacking the matrix metalloproteinase-9 gene reduce renal interstitial fibrosis in obstructive nephropathy. Am J Physiol Renal Physiol, 2010. 299(5): p. F973–82.

